# Full-length isoform sequencing for resolving the molecular basis of Charcot-Marie-Tooth 2A

**DOI:** 10.1101/2023.02.07.526487

**Authors:** Andrew B. Stergachis, Elizabeth E. Blue, Madelyn A Gillentine, Lee-kai Wang, Ulrike Schwarze, Adriana Sedeño Cortés, Jane Ranchalis, Aimee Allworth, Austin E. Bland, Sirisak Chanprasert, Jingheng Chen, Daniel Doherty, Andrew B. Folta, Ian Glass, Martha Horike-Pyne, Alden Y. Huang, Alyna T. Khan, Kathleen A. Leppig, Danny E. Miller, Ghayda Mirzaa, Azma Parhin, Wendy Raskind, Elisabeth A. Rosenthal, Sam Sheppeard, Samuel Strohbehn, Virginia P. Sybert, Thao T. Tran, Mark Wener, University of Washington Center for Mendelian Genomics (UW-CMG), Undiagnosed Diseases Network (UDN), Peter H. Byers, Stanley F. Nelson, Michael J. Bamshad, Katrina M. Dipple, Gail P. Jarvik, Suzanne Hoppins, Fuki M. Hisama

## Abstract

**Objectives:** Transcript sequencing of patient derived samples has been shown to improve the diagnostic yield for solving cases of likely Mendelian disorders, yet the added benefit of full-length long-read transcript sequencing is largely unexplored.

**Methods:** We applied short-read and full-length isoform cDNA sequencing and mitochondrial functional studies to a patient-derived fibroblast cell line from an individual with neuropathy that previously lacked a molecular diagnosis.

**Results:** We identified an intronic homozygous *MFN2* c.600-31T>G variant that disrupts a branch point critical for intron 6 spicing. Full-length long-read isoform cDNA sequencing after treatment with a nonsense-mediated mRNA decay (NMD) inhibitor revealed that this variant creates five distinct altered splicing transcripts. All five altered splicing transcripts have disrupted open reading frames and are subject to NMD. Furthermore, a patient-derived fibroblast line demonstrated abnormal lipid droplet formation, consistent with MFN2 dysfunction. Although correctly spliced full-length *MFN2* transcripts are still produced, this branch point variant results in deficient MFN2 protein levels and autosomal recessive Charcot-Marie-Tooth disease, axonal, type 2A (CMT2A).

**Discussion:** This case highlights the utility of full-length isoform sequencing for characterizing the molecular mechanism of undiagnosed rare diseases and expands our understanding of the genetic basis for CMT2A.

## Introduction

Transcript sequencing is emerging as a powerful clinical tool, and recent studies report that transcript sequencing can increase diagnostic yield by 2-24% versus DNA sequencing alone when evaluating suspected Mendelian disorder cases^1–5^. These studies uniformly use short-read sequencing approaches to identify transcripts with aberrant expression levels or spliced products^6^. However, the full-length transcripts produced by these aberrantly spliced products are not typically evident using short-read sequencing, as exon skipping and alternative splice site usage within multi-intronic genes can create transcripts with potential dominant-negative or loss-of-function impacts. Distinguishing among these full-length transcript outcomes is important for appropriately evaluating conditions whereby distinct phenotypes and inheritance patterns are associated with dominant-negative or loss-of-function variants in the same gene.

Charcot-Marie-Tooth 2A (CMT2A) is the most common subtype of CMT2 and is an axonal peripheral nerve disorder characterized by motor, sensory, or autonomic neuropathy. ∼90% of CMT2A cases follow an autosomal dominant inheritance pattern associated with dominant-negative variants in mitofusin 2 (*MFN2*)^7^. In contrast, autosomal recessive inheritance is associated with biallelic loss-of-function (LOF) *MFN2* variants, which typically do not result in a clinical phenotype in the heterozygous state. Splicing variants in *MFN2* can cause both dominant and recessive forms of CMT2A^8–10^, indicating the need to accurately identify the full-length transcript effect of novel splicing variants.

Recent advances in highly accurate long-read, full-length transcript sequencing has the potential to aid in the evaluation of splice-site variants. We report a patient who was found by a combination of short-read and full-length transcript sequencing to have a homozygous branch point variant in *MFN2* that results in deficient MFN2 protein levels via the creation of five distinct altered transcripts that are all subject in nonsense-mediated decay (NMD).

## Methods

### Exome sequencing and analysis

Quad exome sequencing (proband, unaffected mother, unaffected brother, unaffected paternal-half-brother) was performed on DNA (Baylor College of Medicine) through the Undiagnosed Diseases Network (UDN). In addition to clinical exome analysis, researchers reprocessed the exome data and performed queries focused on UPD(1) and genes previously implicated in CMT and similar disorders.

### Short-read transcript/RNA sequencing and analysis

RNA extraction, library preparation, and short-read sequencing were performed on cultured skin fibroblasts from the proband as previously described^5^. A control dataset of short-read transcript/RNA sequencing from 236 skin fibroblast samples, generated at UCLA, was used to identify RNA expression outliers and aberrant splicing products using OUTRIDER^11^ and IRFinder^12^ respectively.

### Full length isoform long-read transcript sequencing and analysis

Cultured skin fibroblasts were grown in DMEM with 10% FBS and then incubated in the presence of cycloheximide (100μg/ml, Sigma-Aldrich) for 6 hours before RNA extraction using the RNeasy Mini kit (Qiagen). Complementary DNA (cDNA) synthesis was performed following the ISO-Seq protocol, which preserves 3’ and 5’ end information (PacBio, Menlo Park, CA). A PacBio SMRTbell library was constructed using these PCR-amplified full-length cDNA transcripts and sequenced using a Sequel II. Iso-Seq data was processed using the Iso-Seq3 pipeline, mapped to GRCh38, and visualized using IGV.

### Sanger validation

RNA was isolated as above, and cDNA was synthesized with random hexamers and SuperScript™ III reverse transcriptase (Invitrogen). The *MFN2* region of interest was amplified by PCR with a sense primer in exon 5 (5’-GCCATGAGGCCTTTCTCCTT) and an antisense primer in exon 8 (5’- AGACGCTCACTCACCTTGTG). PCR products were separated on 7% polyacrylamide gel. Normal and all abnormal products were excised from the gel and DNA was retrieved by submersion of gel slices in 100μl water at room temperature overnight. Eluted PCR products were amplified using the same primers in exons 5 and 8 and amplicons were subjected to Sanger sequencing.

### Lipid droplet analysis

Skin fibroblast cultures from the proband and an unrelated patient who does not harbor variants in *MFN2* were separately plated on glass bottom dishes (MatTek). After 48 hours of culture, cells were incubated with 0.1μg/ml Mitotracker Red CMX Ros (Molecular Probes), 5mM BODIPY 493/503 (Invitrogen) and 3 drops of NucBlue (Invitrogen) for 15 min at 37°C with 5% CO2. Cells were subsequently moved into complete media for ≥45 minutes, then imaged using a Z-series step size of 0.3μm on a Nikon Ti-E widefield microscope with a 63X NA 1.4 oil objective (Nikon), solid-state light source (Spectra X, Lumencor), and an sCMOS camera (Zyla 5.5 megapixel). Each line was imaged on three separate occasions by a blinded experimenter (n>100 cells per experiment). Images were deconvolved using 7 iterations of 3D Landweber deconvolution. The number and fluorescence intensity of lipid droplets on deconvolved images was quantified using Spot Detection Analysis (Nikon Elements). Maximum intensity projections were generated using ImageJ software (NIH). All quantification was performed by an experimenter blinded to sample identification.

### Standard Protocol Approvals, Registrations, and Patient Consents

This study was approved by the National Institutes of Health (NIH) Institutional Review Board (IRB) (IRB # 15HG0130), and written informed consent was obtained from all participants in the study.

## Results

### Clinical phenotype associated with MFN2 deep intronic variant

We evaluated a 42-year-old woman who initially presented with abnormal “foot-slapping” gait at one year of age that progressed into distal leg weakness requiring a wheelchair by age 8. She underwent spinal fusion and Harrington rod placement for scoliosis in her teens and developed respiratory involvement in her thirties. She had normal cognitive development and no family history of neuromuscular disease. Electromyography and nerve conduction velocity studies at age 2 years revealed distal motor and sensory polyneuropathy, with positive waves and fibrillation. Nerve and muscle biopsy revealed marked denervation atrophy. Neurologic exam at age 42 showed a quadriparetic woman with normal facial strength, hypophonia, severe muscle wasting of arms and legs, and 1-2/5 proximal motor strength and 0/5 distal strength. Sensation was present but reduced to all modalities distally and reflexes were absent throughout.

### Identification of a deep intronic MFN2 variant using short-read transcript sequencing

Initial genetic evaluation revealed paternal uniparental isodisomy of chromosome 1 (UPD[1]), while panel testing for genes associated with neuromuscular disorders was non-diagnostic. She was enrolled into the UDN, where initial exome analysis did not identify a strong candidate variant. Short-read transcript sequencing of RNA isolated from a patient-derived fibroblast line identified *MFN2*, located on chromosome 1, as an expression outlier in comparison to sequencing data from control fibroblast lines (Z-score -6.9) (**Figure 1A**) with about 2-fold lower expression relative to control fibroblasts. In addition, *MFN2* exhibited increased retention of intron 6 (Z-score 8.6) (**Figure 1B**). Re-analysis of the exome data identified a homozygous *MFN2* c.600-31T>G variant within intron 6 that is absent from population databases and is predicted to disrupt the U nucleotide in the yUnAy consensus branch point sequence^13^ (**Figure 1B**).

**Figure 1.**
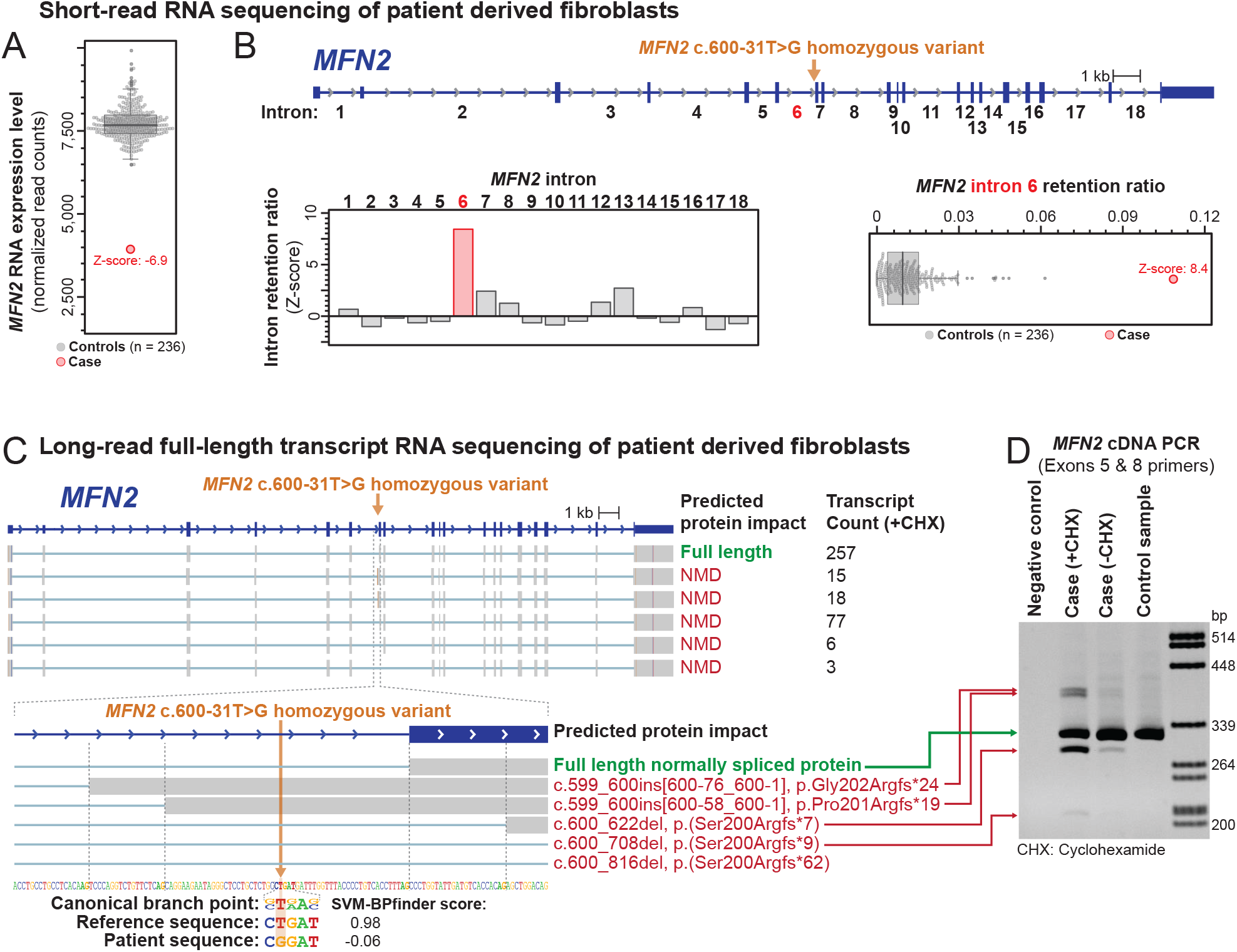
Identification of a homozygous *MFN2* branch point variant that disrupts *MFN2* splicing. (A) Short-read RNA sequencing identified *MFN2* as an expression outlier in this patient’s sample, exhibiting 51% of the RNA expression level seen in control fibroblast samples. (B) Genetic testing of *MFN2* identified a deep intronic homozygous variant in intron 6 of *MFN2*. Short-read RNA sequencing identified that the intron retention ratio of intron 6 of *MFN2* was significantly abnormal compared to controls. (C) Long-read full-length transcript sequencing (ISO-Seq) of this patient’s sample after treatment with the non-sense mediated decay (NMD) inhibitor cycloheximide (CHX) identified 6 major *MFN2* transcripts. The predicted protein impact and transcript count of each are indicated to the right. Inset below shows the alternative splice acceptor sites used for each transcript, as well as the sequence context of the patient’s variant relative to the canonical branch point sequence. (D) DNA agarose gel showing altered spliced products affecting exon 7 of *MFN2* before and after CHX treatment, as well as in a control sample.

### Identification of the splicing impact of an MFN2 branch point variant

Heterozygous LOF *MFN2* variants are not typically associated with disease, and this patient’s MFN2 transcript level was only reduced to 51% of normal (**Figure 1A**). As branch-point variants can induce complex splicing alterations^14^, we performed full-length isoform sequencing (ISO-Seq) to determine the identity of all full-length *MFN2* spliced transcripts. ISO-Seq data from patient-derived cells treated with the NMD inhibitor cycloheximide revealed five distinct altered *MFN2* transcripts that each use a distinct splice acceptor site in lieu of the canonical exon 7 splice acceptor site (**Figure 1C**). Notably, all five altered splicing transcripts have disrupted open reading frames that make them subject to NMD, and none of them are present within control fibroblast cell lines (**Figure 1D**). Overall, these data demonstrate that this branch point variant does not create a significant amount of a stable abnormal protein but does substantially reduce the amount of normal protein.

### An MFN2 branch point variant causes insufficient MFN2 levels

To determine whether this branch point variant results in insufficient MFN2 protein levels, we analyzed patient-derived fibroblast cells for hallmarks of *MFN2* dysfunction. MFN2 is essential for mitochondrial dynamics, and pathogenic *MFN2* variants are associated with diverse mitochondrial phenotypes, including impaired mitochondrial respiration and movement, as well as increased lipid droplet formation^15^. We found that patient-derived fibroblast cells had both increased number and intensity of lipid droplets compared to control cells (**Figure 2**), which is consistent with the idea that this branch point variant results in insufficient functional MFN2 protein.

**Figure 2.**
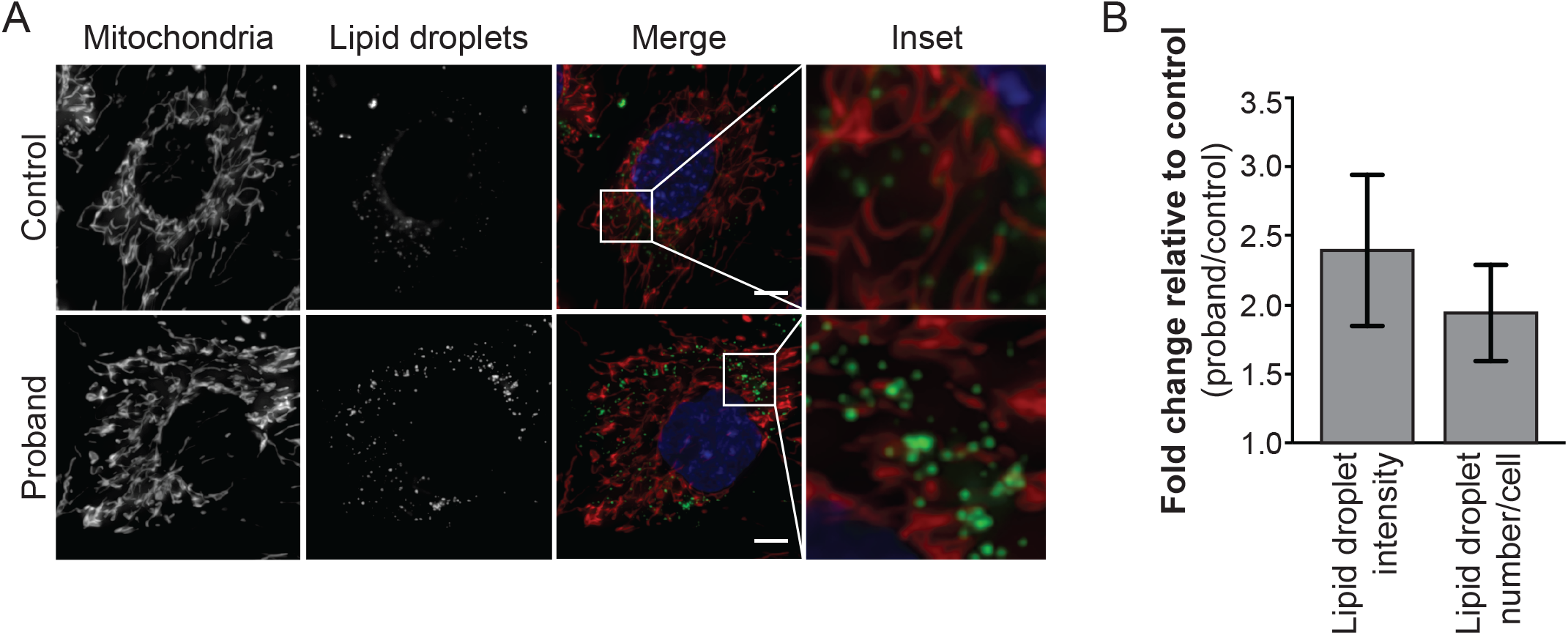
*MFN2* branch point variant results in abnormal lipid droplet formation. (A) Representative images of control and proband fibroblast cells. Mitochondria were labeled with Mitotracker CMXRos, lipid droplets with Bodipy 493/503, and nuclei with NucBlue. Images represent maximum intensity projections. Scale bar = 5 μm. (B) Fold increase of lipid droplet fluorescence intensity and number in proband compared to control.

## Discussion

This study describes a pathogenic intronic branch point variant that alters splicing of intron 6 in both copies of *MFN2* sufficiently to alter the total functional capacity of the encoded protein – expanding our understanding of the molecular basis of CMT2A. Full-length transcript sequencing allowed us to identify and quantitate the outcomes of abnormal intron 6 splicing, exposing that this is a leaky variant that results in some normally spliced transcripts. Our data suggests that the overall lower transcript abundance is sufficient to produce this phenotype. This is consistent with a lack of symptoms in her father, who is heterozygous for this variant, and indicates that this variant results in an autosomal recessive LOF mechanism.

Recent advances in long-read sequencing technology and full-length transcript sequencing have the potential to transform clinical workflows for evaluating patients with unsolved likely Mendelian conditions. This study provides a proof-of-concept for the utility of full-length transcriptome data to identify disease-associated variants and to characterize the mechanism by which these variants cause disease. Further studies are needed to fully evaluate the utility of full-length transcript data in clinical practice.

## Study Funding

A.B.S holds a Career Award for Medical Scientists from the Burroughs Wellcome Fund and is a Pew Biomedical Scholar. This study was supported by NIH grants 1U01HG010233, 1DP5OD029630, R01GM118509 and U01HG007703, in addition to funds from the Collagen Diagnostic Laboratory, University of Washington. Fibroblast data and analysis were partially supported by the California Center for Rare Diseases within the UCLA Institute of Precision Health. Sequence data analysis was supported by the University of Washington Center for Mendelian Genomics (UW-CMG), which was funded by NHGRI grant UM1 HG006493. The content is solely the responsibility of the authors and does not necessarily represent the official views of the National Institutes of Health.

## Disclosure

The authors report no competing interests.

## Notes

### Competing Interest Statement

The authors have declared no competing interest.

